# Postnatal Pulmonary Artery Development from Transcript to Tissue

**DOI:** 10.1101/2025.06.03.657639

**Authors:** Erica L. Schwarz, Abhay B. Ramachandra, Nicola Yeung, Edward P. Manning, Dar Weiss, Jay D. Humphrey

## Abstract

Many congenital conditions and surgical interventions perturb the hemodynamics experienced by proximal pulmonary arteries during early postnatal development, thus leading to differential gene expression and associated changes in vascular structure and function. Among these, pathologic conditions include patent ductus arteriosus, pulmonary atresia and stenosis, and hypoxemia-induced pulmonary hypertension while surgical interventions include the placement of a Blalock-Taussig shunt as well as Glenn, Fontan, and Norwood procedures. Despite the significant morbidity associated with these diverse conditions, there has been little attention directed towards understanding natural postnatal development of pulmonary arteries from both biological and mechanical perspectives. Without such information, we cannot truly understand the phenotype of the affected pulmonary artery, which is fundamental to improving diagnosis, treatment, and prognosis. In this paper, we present novel data from wild-type mice that document normal postnatal changes in select gene expression, vascular wall composition, and biomechanical properties of proximal pulmonary arteries. These findings enabled the establishment of a novel, data-informed computational model of pulmonary artery development capable of simulating outcomes in response to perturbations in the pulmonary artery hemodynamic environment.

## 1 Introduction

Pulmonary arteries serve the vital function of conducting blood to the lungs for gas exchange. Normal postnatal development of these arteries can be disrupted by diverse congenital defects, early onset diseases, or early clinical interventions, including staged surgical palliation. Examples of maldevelopment of pulmonary arteries due to congenital conditions include vessel hypertrophy in left hypoplastic heart syndrome (*1*), vessel atresia in ventricular septal defects (*2*), altered medial composition and wall thickness in patent ductus arteriosus (*3, 4*), and aberrant remodeling in pulmonary stenosis (*5*), among others. Although beneficial overall, surgical interventions such as placement of a Blalock-Taussig shunt and performance of Norwood, Glenn, and Fontan procedures can further compromise normal vessel development (*6–9*). Many studies have used computational analyses of the hemodynamics to understand altered pulmonary blood pressure and flow under particular conditions (*10–14*), but none have sought to model, and thus understand quantitatively, the associated developmental changes in gene expression, mural composition and structure, and biomechanical properties that ultimately dictate pulmonary artery function and clinical outcomes. Toward this end, there is a need to understand better the time-course of normal prenatal and postnatal development of pulmonary artery structure and function.

Although greater attention has traditionally been placed on small (intra-parenchymal) pulmonary arteries because of their central role in pulmonary hypertension (*15*), it is becoming increasingly evident that the structure and function of large (extra-parenchymal) pulmonary arteries also play important roles in ventilation-perfusion (*16–18*). Unlike conduit arteries in the systemic circulation, healthy proximal pulmonary arteries typically experience a reduction in blood pressure at birth with subsequent near maintenance of pressure despite monotonically increasing cardiac output and thus blood flow. As a result, normal proximal pulmonary arteries develop a unique biomechanical phenotype in maturity that includes both moderate elastic energy storage capability characteristic of a conduit artery and marked contractile capacity characteristic of a muscular artery (*19*).

Previous studies have attempted to develop computational models of proximal arteries during postnatal development, but have focused on the aorta (*20, 21*). These models assume that homeostatic intramural and flow-induced shear stress targets evolve during development, yet evidence suggests that certain homeostatic targets may be maintained throughout embryonic and postnatal development, with particular evidence that wall shear stress (WSS) is closely maintained at near-mature values during late embryonic development (*22*) and, with the exception of transient deviations, tends to change little during postnatal development (*21, 23*). Therefore, there is still a need for a computational model that accounts for the dynamic evolution of the postnatally developing pulmonary artery while considering certain homeostatic targets.

Herein, we report new multi-modality data of the primary branch pulmonary artery throughout postnatal development in normal mice at five developmental ages from just after birth to maturity. We then use these data to establish a computational model of proximal pulmonary artery development rooted in the constrained mixture theory of tissue growth (changes in mass) and remodeling (changes in structure). We also demonstrate the ability of this framework to simulate known outcomes of pulmonary artery growth and remodeling (G&R) in response to hemodynamic perturbation, both during development and in maturity.

## 2 Methods

### 2.1 Animals

Motivated by data on time-course changes in biomechanical structure and function during postnatal development of the thoracic aorta in mice (*23*), we studied development of the proximal pulmonary artery in male C57BL/6J mice at postnatal days P2, P10, P21 (weaning), P42, and P84 (well into maturity) following an Institutional Animal Care and Use Committee of Yale University approved procedure for euthanization (CO2 exposure followed by decapitation at P2 and P10, CO2 asphyxiation for P21-P84) and tissue harvest. The right branch pulmonary artery (RPA) was isolated via a midline sternotomy and gently excised. Following ARRIVE Guidelines (*24*), excised vessels were randomly divided into three cohorts: those for bulk RNA sequencing (*n* = 5 at P2, P10, P21, P42, P84), those for biomechanical testing (*n* = 3−5 at P2, P10, P21, P42, P84), and those for histological analysis (*n* = 4−6 at P2, P10, P21, P42, P84).

### 2.2 RNA-Sequencing

To promote reproducibility, we used a previously established protocol (*25*). Immediately upon euthanasia, RNA was isolated from the extralobar pulmonary arteries using an RNeasyMini Kit (Qiagen) according to the manufacturer’s specifications. Quality control measures ensured that samples were not degraded, with RNA integrity (RIN) ¿ 7. RNA libraries were prepared with polyA selection. Whole-transcriptome sequencing was performed using a NovaSeq 6000 System (Illumina, Inc.) by The Yale Center for Genome Analysis. Sequenced reads were imported into CLC Genomics Workbench V23 (Qiagen) and, following quality control, reads were trimmed and aligned/mapped to a *Mus musculus* reference genome. Reads were then automatically processed using log counts per million (CPM) transformation with trimmed mean of M (TMM) adjustment. Although we collected full transcriptional information, herein we focus on the transcripts that relate directly to measurable changes in biomechanical constituents, *Eln, Col1a1, Col3a1, Col5a1*, and *Pcna*.

### 2.3 Biomechanical Phenotyping

To increase rigor and reproducibility, we followed standard procedures established in our laboratory for murine vessels (*19, 26*). Upon euthanasia, the RPA was isolated by gentle excision, cleaned of excess perivascular tissue, cannulated on custom glass cannulae, and placed within a specimen bath containing a Hank’s buffered physiologic solution at room temperature, which ensures a passive mechanical behavior. Using a custom computer-controlled test device (*27*), we performed standard preconditioning, then subjected the vessels to a series of seven cyclic pressure-distension and axial force-extension protocols that generated pressure, diameter, axial force, and axial length data. The energetically favorable *in vivo* axial stretch was estimated as that value at which axial force changed little upon cyclic pressurization. Vessels were subjected to pressures up to which asymptotic behavior was observed in the pressure-diameter curves (20 mmHg at P2, 25 mmHg at P10, 30 mmHg at P21, and 40 mmHg at P42 and P84). The axial stretch was maintained constant at 95%, 100%, and 105% of the in vivo value. During the force-length tests, vessels were subjected to axial forces up to the maximum value achieved during the 105% stretch pressure-diameter test while maintaining the luminal pressure fixed at four different values at or below the maximum testing pressure.

### 2.4 Histology

Following mechanical testing, all samples were fixed in 10% neutral buffered formalin and stored in 70% ethanol at 4^°^C. Samples were embedded in paraffin, sectioned, and stained with Movat pentachrome (MOVAT) by a Yale histology core. Under bright-field illumination, MOVAT reveals elastin as black, collagen as grey-yellow, glycosaminoglycans (GAGs) as blue, cytoplasm as pink, and, if present, fibrin as dark red. Sections were imaged with an Olympus BX/51 microscope and an Olympus DP70 digital camera at 40X magnification. Complete cross-sections were obtained by stitching together sub-images with the Image Composite Editor software (Microsoft Research). The stitched images were subsequently analyzed using custom MATLAB scripts (*28*) that included background subtraction and pixel-based thresholding. Three sections (technical replicates) were analyzed per vessel (five biological replicates), thus yielding 15 sections per age.

Area fractions for elastin and cytoplasm were computed as the ratio of pixels corresponding to a stain divided by the total number of pixels in the image. RNA sequencing showed high levels of proteoglycan expression (*Acan* and *Vcan*) at neonatal ages that exponentially decayed to low levels at adult ages (Fig. S1). This was in agreement with previous observations of negligible GAG content in the adult pulmonary artery (*19*) which was also observed in this study (Fig. 1). Despite the high expression of *Acan* and *Vcan* at P2 and P10, neontal histology stains did not reveal distinct areas of GAG accumulation at these ages (Fig. 1). This may suggest that GAGs are diffusely mixed with collagen in the extracellular matrix during the neonatal period. To account for this, the remaining pixels not categorized as elastin or cytoplasm in the histological sections are considered to be a combination of collagen and GAGs, with the area fraction of this constituent aggregate computed as 1 - area fraction of elastin plus cytoplasm. Given the diffuse GAGs and their negligible presence in maturity, we model this mixture according to collagen turnover and load-bearing properties.

**Figure 1.**
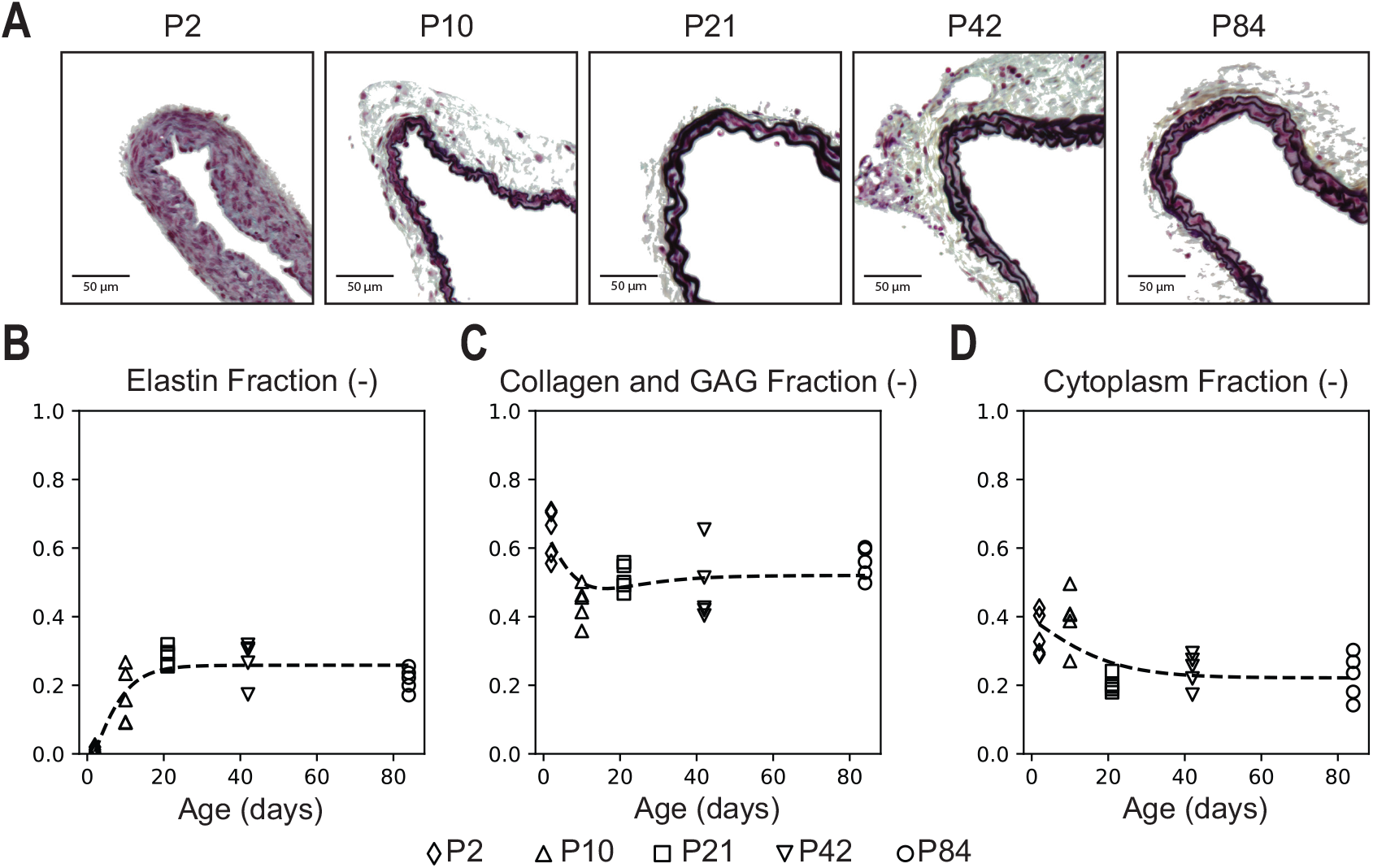
Histological analysis of RPA wall composition during postnatal development. Symbols represent data at respective ages; dashed trendlines are shown for visualization. (A) Movat pentachrome (MOVAT) stained sections with background subtraction for representative RPAs at P2, P10, P21, P42, and P84. This staining allowed quantification of elastin (black), collagen (grey/yellow), GAGs (blue), and cytoplasm (pink). Scale bars indicate 50 μm. (B) The area fractions for elastin and cytoplasm were computed based on n=5 sections for each stain at each age, with the remaining collagen plus GAG fraction calculated as 1 - area fraction of elastin and cytoplasm.

### 2.5 Constitutive Modeling

We model the evolving vessel using a constrained mixture theory (*29*), whereby the mechanical behavior of individual constituents determine the stress response of the overall vessel such that the Cauchy stress, **σ**, of the tissue is

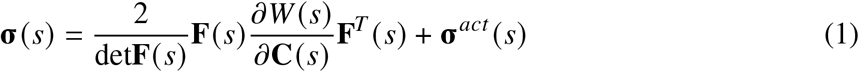

where **F**(*s*) is the deformation gradient tensor tensor at G&R time *s*, **C**(*s*) = **F**^*T*^ (*s*)**F**(*s*) is the right Cauchy-Green tensor, **σ**^*act*^ (*s*) is the active stress contribution, and *W* (*s*) is the mixture-level strain energy for passive behavior at time *s*.

In constrained mixture theory, a simple rule of mixtures relation is adopted for *W* (*s*), which conceptually has the form *W* (*s*) = ∑ _*α*_ *ϕ*^*α*^ (*s*)*W*^*α*^ (*s*) where *W*^*α*^ (*s*) is the constituent-specific strain energy at time *s* and *ϕ*^*α*^ (*s*) is the current constituent mass fraction at time *s*. We define as *ϕ*^*α*^ (*s*) = *ρ*^*α*^ (*s*)/*ρ*(*s*) where *ρ*^*α*^ (*s*) is the apparent mass density of constituent *α* at time *s* and *ρ*(*s*) is the mass density of the mixture as a whole at time *s*.

Consistent with a mixture-based mass balance relation, we model the potentially evolving constituent-specific apparent mass density as

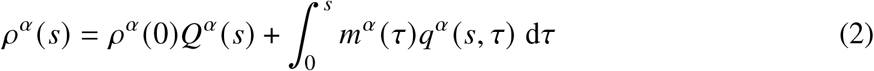

where *m*^*α*^ (*τ*) *>* 0 is the mass density production at G&R time *τ* and *q*^*α*^ (*s, τ*) ∈ [0, 1] is the fraction of constituent *α* deposited at time *τ* that survives until time *s. ρ*^*α*^ (0) is the mass density at the initial G&R time (taken herein to be at P2), while *Q*^*α*^ (*s*) ∈ [0, 1] is the fraction of this initial constituent cohort that survives until time *s*. The constituent strain energy then is assumed to be (*30*)

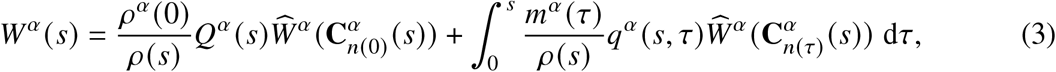

where 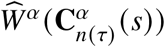 is a mass-averaged, constituent-specific strain energy function and 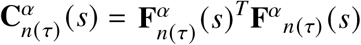 is the constituent-specific right Cauchy-Green tensor that arises from the constituent-specific deformation gradient 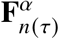 that specifies the deformation of a constituent at time *s* that was deposited at an earlier time *τ* ∈ [0, *s*]. It can be shown that 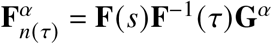, where **F**(*s*)**F**^−1^ (*τ*) represents the deformation of the mixture since deposition at time *τ* and **G**^*α*^ accounts for cell-mediated deposition of structural constituents having a preferred prestretch (*30*).

Typically, we model mass density production *m*^*α*^ (*τ*) via a basal value that can be modulated in response to cellular sensing of mechanical and chemical signals. Therefore,

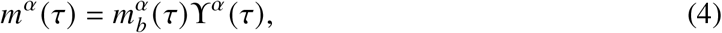

where 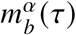 is a basal production rate and ϒ^*α*^ (*τ*) is a chemo-mechanical stimulus function. Because most matrix degradation and cell death can be modeled using first-order kinetics, we let the survival function have the form

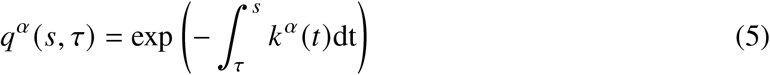

where *k*^*α*^ (*t*) is a rate parameter that can also be modulated in response to external stimuli. Below, we provide the specific forms of the production, removal, and strain energy for the primary load-bearing vascular constituents: elastic fibers organized into laminae, families of oriented collagen fibers plus associated GAGs, and smooth muscle cells (SMCs). Note that these relations seek to capture G&R. By constrast, simple geometric relations capture homeostatic relations for radius (*a* → *ε*^1/3^*a*_***h***_) and wall thickness (***h*** → *γε*^1/3^***h***_***h***_) where *ε* is a fold change in flow and *γ* is a fold change in pressure, with subscript ***h*** denoting homeostatic values (*31*).

### 2.6 Constituent Production and Removal

#### 2.6.1 Collagen and GAGs

Deposition of vascular collagen is mediated by mechanical stimuli (*32, 33*). Here, we define the stimulus function ϒ^*c*^ (*τ*) as

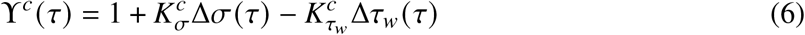

where 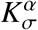 and 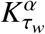 are gain-like parameters that modulate production in response to deviations in pressure- and axial force-induced intramural Cauchy stress and flow-induced WSS from homeostatic values. We let

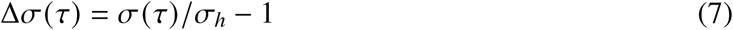

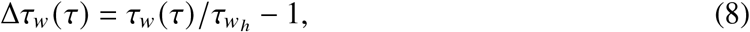

where *σ*(*τ*) is the current first invariant of the 3D Cauchy stress, *τ*_*w*_ (*τ*) is the current WSS, and *σ*_***h***_and 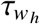 are the homeostatic target values of each in maturity (*31*).

Previous studies have noted that the natural turnover rate of collagen is extremely high *in utero*, but it declines throughout postnatal development to its final, low homeostatic value in maturity (*34, 35*). To account for this behavior, we let the removal rate start at a high value at P2 and then exponentially decay to its value in maturity such that

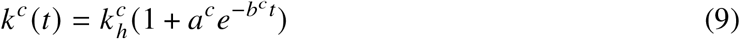

where *a*^*c*^ and *b*^*c*^ are constants and 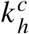 is the homeostatic removal rate in maturity.

It is convenient to similarly parameterize the basal mass production value using this rate such that

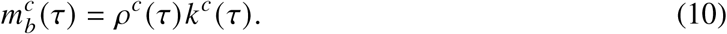

Note that using Eqn. 9 and Eqn. 10 in Eqn. 4 and Eqn. 5 accounts for the much higher turnover observed in development while recovering the well-known behavior of collagen turnover in maturity. In addition, it accounts for the hypothesized removal of neonatal GAGs in the collagen-GAG mixture during the neonatal period.

#### 2.6.2 SMCs

A recent comparison across the great vessels suggests that SMCs seek to achieve a preferred density in maturity (*36*). That is, if the density is too low, SMCs proliferate; if the density is too high, SMCs apoptose. To model this behavior, we define the value:

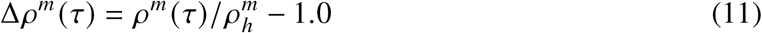

where 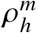 is the homeostatic mass density of SMCs. It has also been noted that neonatal SMCs appear to have much higher rates of replication than adult SMCs (*35*). Therefore, we also define an exponentially decaying basal production rate

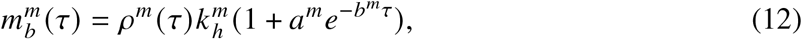

where *a*^*m*^ and *b*^*m*^ are constants and 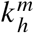 is the homeostatic removal rate in maturity. We then define ϒ^*m*^ (*τ*) and *k*^*m*^ (*t*) in terms of this value such that

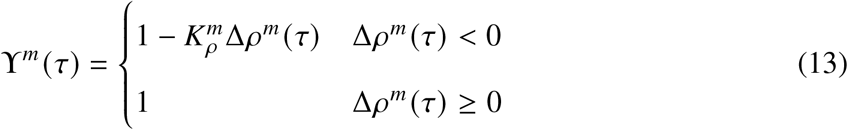

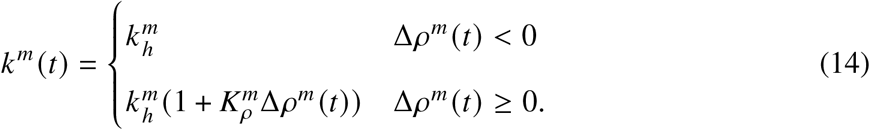

such that, in maturity, there is no change in overall SMC mass density in the absence of production or removal stimuli.

#### 2.6.3 Elastin

Vascular elastin is produced primarily by SMCs. We model the production of elastin as arising from a metabolic stimulus function and the current mass density of SMCs in the tissue such that

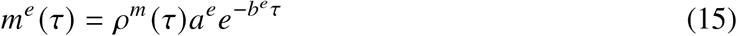

where *a*^*e*^ and *b*^*e*^ are constants. We assume negligible removal of functional elastin once it is deposited, in agreement with observations that elastin has a half-life on the order of 50 years *in vivo* (*37*). Heightened degradation can occur in certain diseases, but this is not considered here.

### 2.7 Constituent Strain Energy Forms

#### 2.7.1 Collagen and GAGs

We model the nonlinear mechanical behavior of collagen fibers with a Fung-type exponential relation having a preferential fiber direction. Let

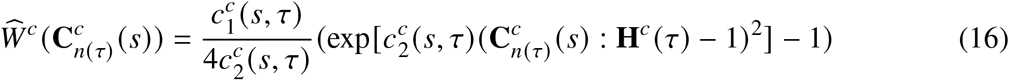

where **H**^*c*^ (*τ*) = **h**^*c*^ (*τ*) ⊗ **h**^*c*^ (*τ*) is the fiber direction tensor with **h**^*c*^ (*τ*) the orientation of the collagen with respect to the axial direction at G&R time *τ*. Note that 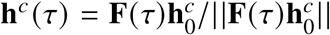 with 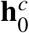 the fiber direction of constituent *α* in the reference configuration. The deposition tensor is given by

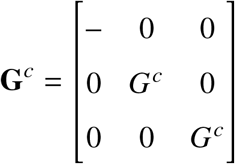

where the scalar function *G*^*c*^ represents the prestretch of the collagen fibers that occurs at deposition due to SMC actomyosin activity. The hyphen in the prestretch tensor represents the lack of collagen fiber orientation in the radial direction.

Previous studies have shown that collagen undergoes a maturation process after deposition (*38*).

We model a similar maturation process in the 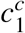 parameter, representing a maturation of cross-link strength. We represent this maturation as a sigmoid

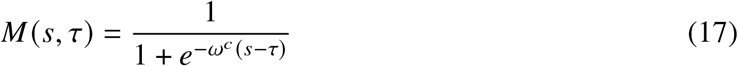

where *M* (*s, τ*) ∈ [0.5, 1] ∀ *s* ≥ *τ* is the maturation value and *ω*^*c*^ is the maturation rate of the collagen. As the current time, *s*, continues to increase past the time of deposition, *τ*, the material maturation, *M* (*s, τ*), will approach 1. This maturation process also helps account for the GAG content during the noeonatal period by ensuring that this collagen and GAG mixture is primarily immature with lower load-bearing capabilities at early ages.

We also assume that as collagen is degraded, the strength of its cross linking also declines.

Therefore, we mediate the values of 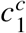 and 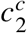 for collagen as a function of the survival fraction such that

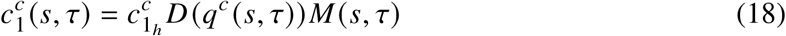

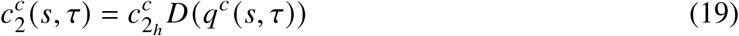

where 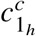 and 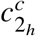 are the maximum values of 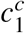 and 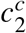, respectively. *D* (*q*^*c*^ (*s, τ*)) ∈ [0, 1] ∀ *q*^*c*^ (*s, τ*) represents a sigmoidal material degradation such that

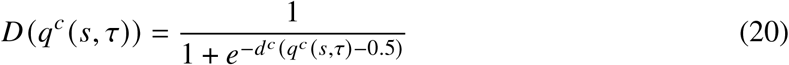

where *d*^*c*^ is a degradation rate. This relationship enforces a 50% decline in cross-linking strength when the collagen family is 50% degraded.

#### 2.7.2 SMCs

We model smooth muscle cell contractility via

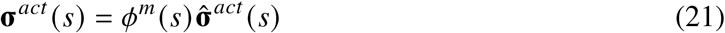

where

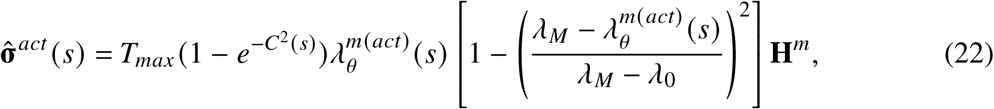

where, **H**^*m*^ = **h**^*m*^ ⊗ **h**^*m*^ and *T*_*max*_ (*s*) is the maximum active stress-generating capacity of the smooth muscle at time *s*. **h**^*m*^ is the fiber direction of the SMCs and is oriented in the circumferential direction of the vessel. *C* (*s*) accounts for the ratio of vasoconstrictors to vasodilators via

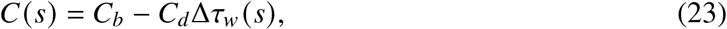

where *C*_*b*_ is a basal ratio and *C*_*d*_ is a scaling factor that accounts for deviations in WSS from homeostatic. 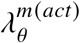 is the circumferential stretch used to calculate active stress, defined as

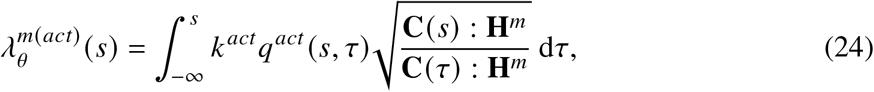

where we let

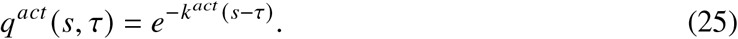

This function represents relaxation of vasomotor tone over time.

#### 2.7.3 Elastin

We model the mechanical behavior of the elastin-dominated part of the extracellular matrix as a neo-Hookean material of the form

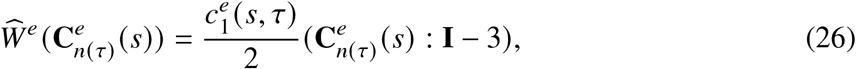

where 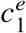 is an evolving shear modulus. The structure of elastin is typically understood to support significant stress in the axial-circumferential direction, but to have relatively sparse connections in the radial direction. Therefore, we use the deposition tensor to capture possible anisotropic responses in the radial direction, namely

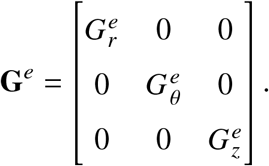

It has been observed that the material constant 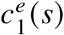 increases during neonatal development (*23*). This is speculated to be due to cross-linking and compaction of the elastin after deposition (*39*). Therefore, we parameterize 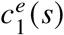 using the sigmoidal relationship

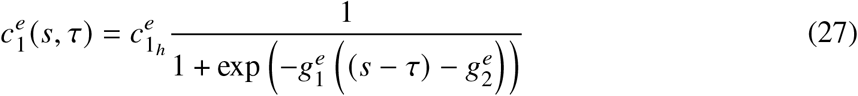

where 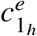 is the elastin material parameter in maturity. Values of 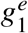 and 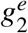 are given below and enforce elastin reaching 50% of its mature stiffness at P21, which is supported by previously reported values for the aorta (*23*).

## 3 Results

### 3.1 Histology

Representative MOVAT sections are shown in Fig. 1 for the RPA at P2, P10, P21, P42, and P84. Elastic fibers in the form of laminae rise from an ∼0% area fraction at P2 to an ∼22% area fraction in maturity at P84. The cytoplasm area fraction, which is typically understood to be proportional to the area fraction of the SMCs, began at ∼35% at P2 before decreasing to ∼23% at P84, inline with previous observations of cell nuclei area fraction in mature arteries (*36*). Together, the accumulation of elastin and reduction in cytoplasm resulted in the area fraction of the collagen and GAG mixture decreasing from ∼64% at P2 to ∼56% at P84 (Table S1). The values of all three area fractions at P84 are consistent with previously measured values in maturity (*19*,*40*). Note, too, the development of the elastin laminae within the medial layer results in two musculo-elastic layers in maturity (Fig. 1).

### 3.2 Loading Conditions

As noted, we assume that mean arterial pressure remains nearly constant throughout the normal postnatal developmental period following closure of the ductus arteriosus. To characterize the hemodynamic environment of the RPA, we also calculate changes in both flow and axial force over time. To estimate RPA flow, we first take the measured body mass of each mouse and use the allometric scaling relationship Cardiac output (ml/min) = 1.0767 · [mass (g)] as reported in (*39*). To calculate RPA flow in particular, we use a reported RPA to left pulmonary artery (LPA) flow ratio 68%:32% (*41*). This yields the time-resolved RPA flow shown in Fig. 2. Given the estimated RPA flow and the measured inner diameter (Fig. 3), we can estimate the evolving mean WSS using the formula *τ*_*w*_ = 4*µQ*/(*πa*^3^) where *µ* is the blood viscosity, *Q* is the flow rate, and *a* is the inner radius. We use *µ* = 4 mPa · s. The resulting trend in *τ*_*w*_ is shown in Fig. 2; it rises from an average value of ∼6 Pa at P2 to ∼9 Pa in maturity. The latter is in agreement with previous estimates of *τ*_*w*_ of the RPA *in utero* and in maturity (*22*). Finally, we show the estimated axial force needed to achieve the *in vivo* axial stretch of each individual vessel; it increases 10-fold throughout development to its mature value of ∼4 mN.

**Figure 2.**
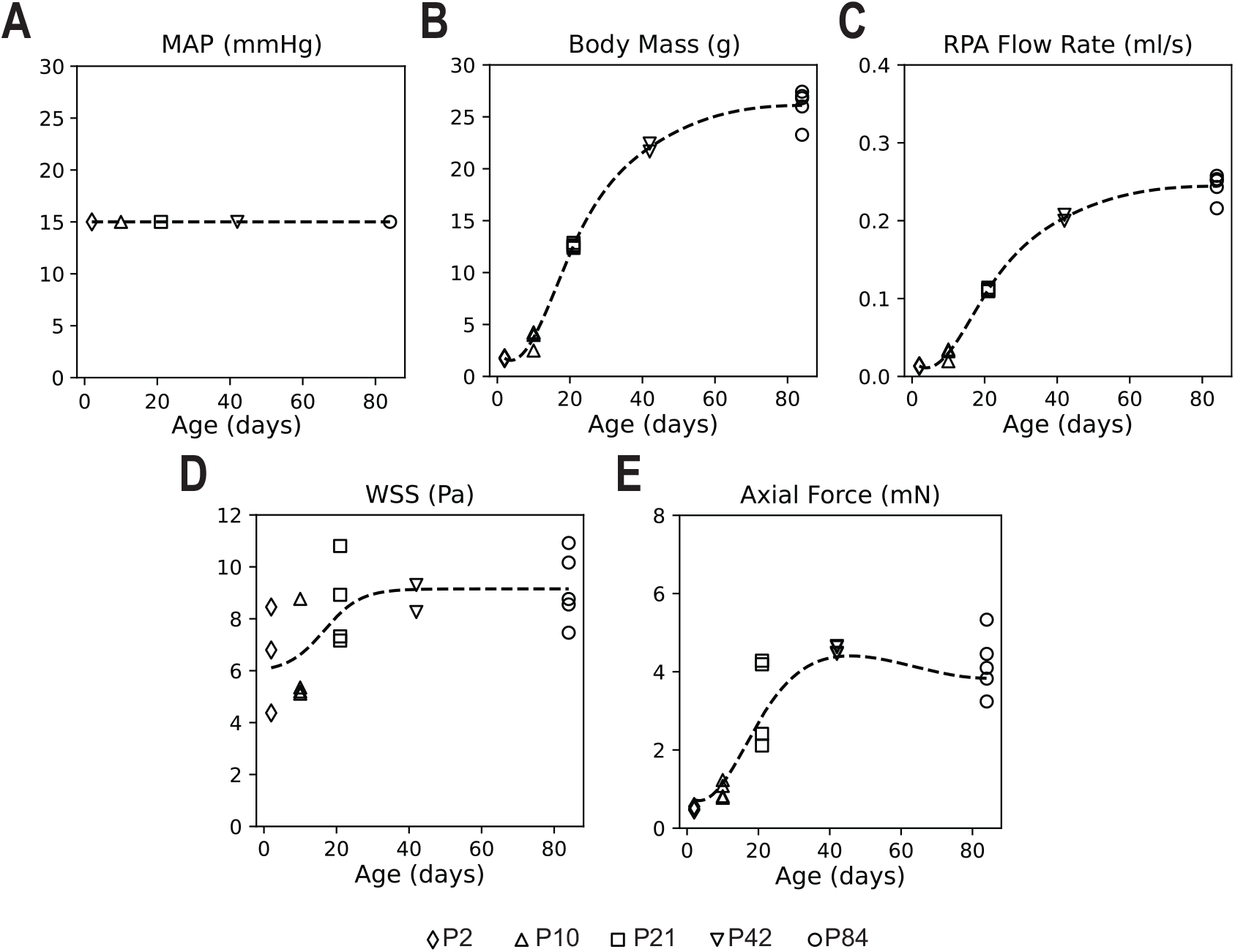
Hemodynamic metrics and loading conditions in the RPA during postnatal development. Symbols represent data at respective ages; dashed trendlines are for visualization. (A) Mean arterial pressure (MAP) in the proximal pulmonary arteries is assumed to drop rapidly after birth from ∼30 mmHg (*52*) and thereafter remain nearly constant at 15 mmHg from P2 to P84. (B) Body mass at each age. (C) RPA flow rate was calculated allometrically using the relationship Cardiac output (ml/min) = 1.0767 · [mass (g)] (*39*). The ratio of RPA to LPA flow was assumed to be constant at 68%:32% (*41*); trendline is used as the inflow boundary condition in the simulation framework. (D) WSS is calculated from RPA flow rate and *in vivo* inner diameter reported in Fig. 3. (E) Axial force that yields the individual *in vivo* axial stretches reported in Fig. 3; trendline is used as the force boundary condition in the simulation framework.

**Figure 3.**
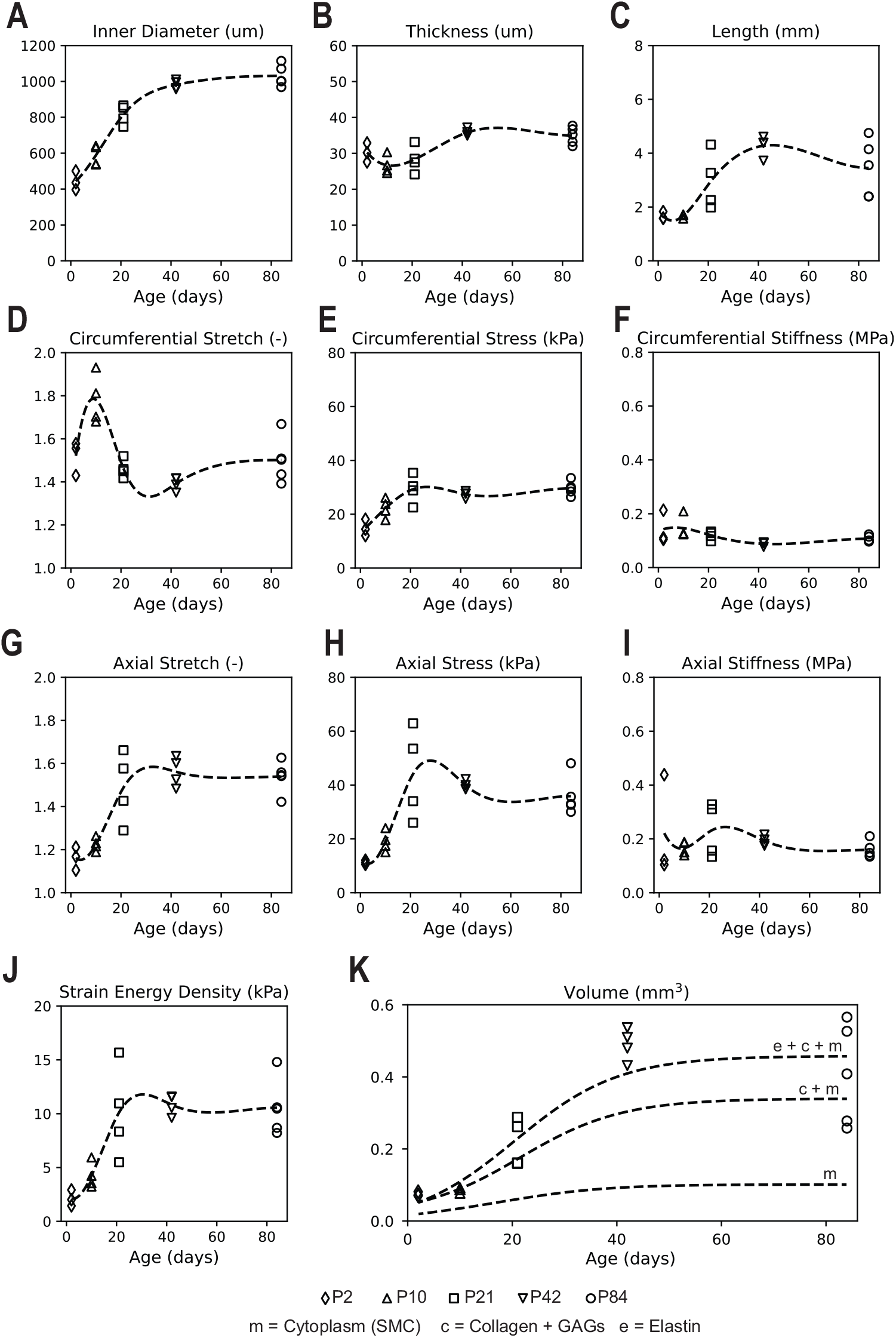
Morphologic and biomechanical data for the RPA during postnatal development. Symbols represent data at respective ages; dashed trendlines are for visualization. Loaded dimensions (A, B, C), biaxial stretches (D, G), stresses (E, H), stiffnesses (F, I), and strain energy density (J) are reported at a mean arterial pressure of 15 mmHg and individual values of the *in-vivo* axial stretch. (K) Total vessel volume is calculated from diameter, thickness, and length. Constituent volumes are calculated by multiplying the mean constituent area fractions reported in Fig. 1 by the total vessel volume and using a sigmoidal fit. Cumulative constituent volumes are overlaid on the volume plot as specified by the figure legend.

### 3.3 Biomechanics

RPAs undergo rapid development after P2. Morphologic and biomechanical values are reported at a mean arterial pressure of 15 mmHg and individual values of *in vivo* axial stretch (Fig. 3, Table S2). The inner radius rises from ∼120 µm at P2 to ∼500 µm at P42 and stays nearly steady thereafter into maturity at P84. Unlike the radius, the loaded thickness of the wall remains relatively constant throughout development at ∼30 µm. Given the constant luminal pressure, this results in an increasing circumferential stress over the first 3 weeks. Axial stress similarly increases over the first 3 weeks, but then reduces slightly to its mature value. By contrast, circumferential and axial material stiffness remain relatively constant throughout postnatal development. Using the morphologic measurements, we can estimate the individual constituent volumes by multiplying vessel volume by mean area fractions reported in Fig. 1, assuming that area fraction is relatively constant along vessel length.

### 3.4 RNA-Sequencing and Turnover

RNA-sequencing yields time-resolved trends in the expression of the elastic fiber precursor gene, *Eln*, the fibrillar collagen gene families *Col1a1, Col3a1*, and *Col5a1*, and the cell proliferation gene, *Pcna* (proliferating cell nuclear antigen). See Fig. S1. We use these trends to calculate data-informed drivers of vascular development. Generally, we postulate that the rate of elastic fiber and collagen fiber production are proportional to the expression of *Eln* and *Col1a1* + *Col3a1* + *Col5a1*, respectively, as introduced in (*39*). In addition, we use *Pcna* to parameterize the replication rate of SMCs throughout development.

For collagen, this can be formalized using the relationship

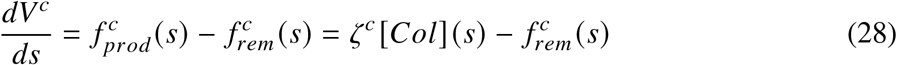

where *dV* ^*c*^/*ds* is the rate of change in collagen volume (Fig. S2) with 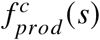 the production rate, 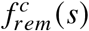 the removal rate, *ζ* ^*c*^ a parameter that describes how production rate scales to gene expression, and [*Col*] (*s*) = [*Col*1*a*1](*s*) + [*Col*3*a*1](*s*) + [*Col*5*a*1](*s*). Generally, we expect the amount of collagen in the pulmonary arteries to be constant in maturity. In addition, previous literature has established that the half-life of aortic collagen is approximately 60-70 days in mature rats (*42*); although for a different species and vascular segment, this range provides an order of magnitude estimate of vascular collagen turnover in maturity. We found that a collagen half-life of 40 days in maturity best fit the mouse pulmonary artery data. We can use these relationships to define, at the homeostatic value (i.e. *dV* ^*c*^/*ds* = 0),

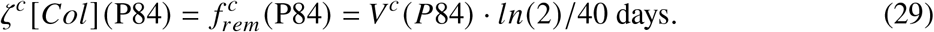

which determines the constant *ζ* ^*c*^. Given the change in collagen volume over time (Fig. S2), we can calculate the time-resolved expressions of 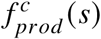 and 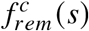. We use this to solve for the relative removal rate, 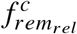 (which is equivalent to *k*^*c*^ (*t*) in Eqn. 9) as shown in Fig. 4 and to parameterize our simulation framework as defined in Section 2.6.1.

**Figure 4.**
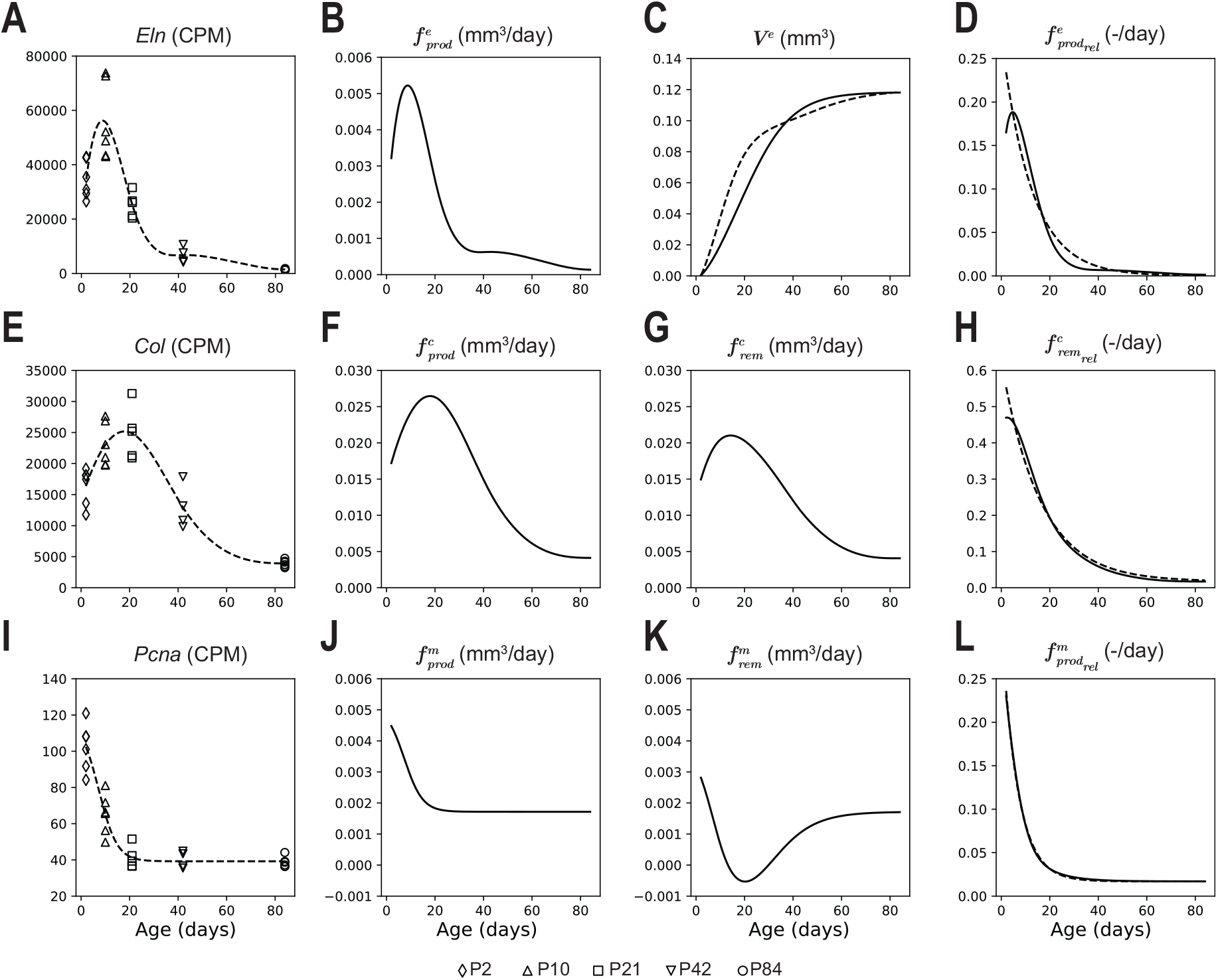
RNA-sequencing and volume turnover data for elastin (*e*), collagen (*c*), and SMCs (*m*). (A, E, I) Gene expression changes for elastin (*Eln*), fibrillar collagen (*Col1a1* + *Col3a1* + *Col5a1*), and cell proliferation (*Pcna*) during postnatal development into maturity; symbols represent data at respective ages. (B, F, J) Production rate of elastin, fibrillar collagen, and SMCs as calculated from RNA-sequencing data. (C) Elastin volume over time calculated from RNA-sequencing data and calculated from the histological and morphologic data. (C, K) Removal rate of fibrillar collagen and SMCs calculated from RNA-sequencing, histological, and morphologic data. Note that the calculation of SMC removal results in a negative removal rate. This can be interpreted as a low, approaching zero, removal rate rather than a truly negative rate. (D) Relative production rate of elastin with respect to current SMC volume; dashed line represents fitted exponential decay used to parameterize constituent production 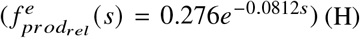 . Relative removal rate of collagen with respect to current collagen volume; dashed line represents fitted exponential decay used to parameterize constituent production and removal 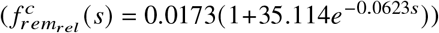. (L) Relative production rate of SMCs with respect to current SMC volume; dashed line represents fitted exponential decay used to parameterize constituent production and removal 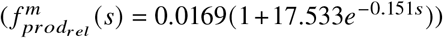 . Note that this calculation of relative SMC production predicts a replication rate of 1.69%/day in maturity, similar to the observed replication rate of ∼2% in mature rat pulmonary arteries (*43*).

Similarly, we assume that the production of elastin is proportional to the expression of *Eln*. However, we assume that there is negligible removal of elastin based on previous studies (*37*). This yields

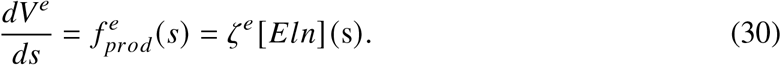

We divide 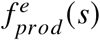 by the current volume of cytoplasm (as a surrogate for SMC volume) to get the production of elastin relative to the current amount of SMCs, 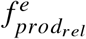, and use this to parameterize our simulation framework as defined in Eqn. 15 in Section 2.6.3. As shown in Fig. 4D, calculating the rate of change of the constituent volume directly from gene expression gives an excellent match to the observed volume changes estimated from the histological and morphometric data.

For SMCs,

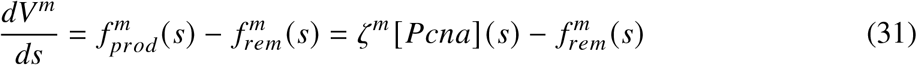

where *dV* ^*m*^/*ds* is the rate of change in SMC volume, 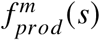 is the production rate, 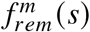 is the removal rate, and *ζ* ^*m*^ is a parameter that describes how production rate scales to the gene expression of [*Pcna*] (*s*). Previous studies have quantified SMC replication rates in neonatal rats, with a measured replication rate at P2 of 23% (*43*). Assuming the replication rate is similar in neonatal mice, we can define

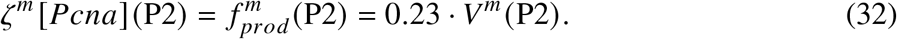

which determines *ζ* ^*m*^. Given the change in cytoplasm volume over time (Fig. S2), we can calculate the time-resolved expressions of 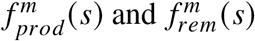. We use this to solve for the relative production rate, 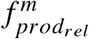, as shown in Fig. 4 and to parameterize our simulation framework as defined in Section 2.6.2. Note that this calculation of relative SMC production predicts a replication rate of 1.69%/day at P84, which agrees with the previously estimated replication rate of ∼2% in mature rat pulmonary arteries (*43*).

### 3.5 Modeling Development

Turnover rates of the major vascular constituents are calculated based on the data defined above. The remaining material parameters were tuned to match the measured, time-resolved biomechanical properties as reported in Section 3.3 based on the loading conditions defined in Section 3.2 (Table S3, Table S4). Linear momentum balance dictates that for a thin-walled cylinder *σ*_*rr*_ ≈ −*P*/2, *σ*_*θθ*_ ≈ *Pa*/***h***, and *σ*_*zz*_ ≈ *f* /[*π****h*** (2*a* + ***h***)], where *a* is inner radius, ***h*** is wall thickness, *f* is axial force, and *P* is transmural pressure. At each time, the mechanical equilibrium solution satisfies the constitutive equations for the prescribed boundary conditions. At P84, we modified the axial boundary condition from a force condition to a displacement condition where the axial length of the pulmonary artery remained constant. This is in line with previous simulations of adult arterial vessels and reflects the limited changes in pulmonary axial length after maturity (*44, 45*). The resulting framework captures the behavior of the RPA during postnatal development, including evolving composition, geometry, stresses, stiffnesses, and strain energy (Fig. 5). Note, in particular, that the model captures both the monotonic increase in radius and nonmonotonic changes in wall thickness and the increase, then decrease back down to homeostatic values, in both axial stress and stiffness as well as trends in all other metrics of interest.

**Figure 5.**
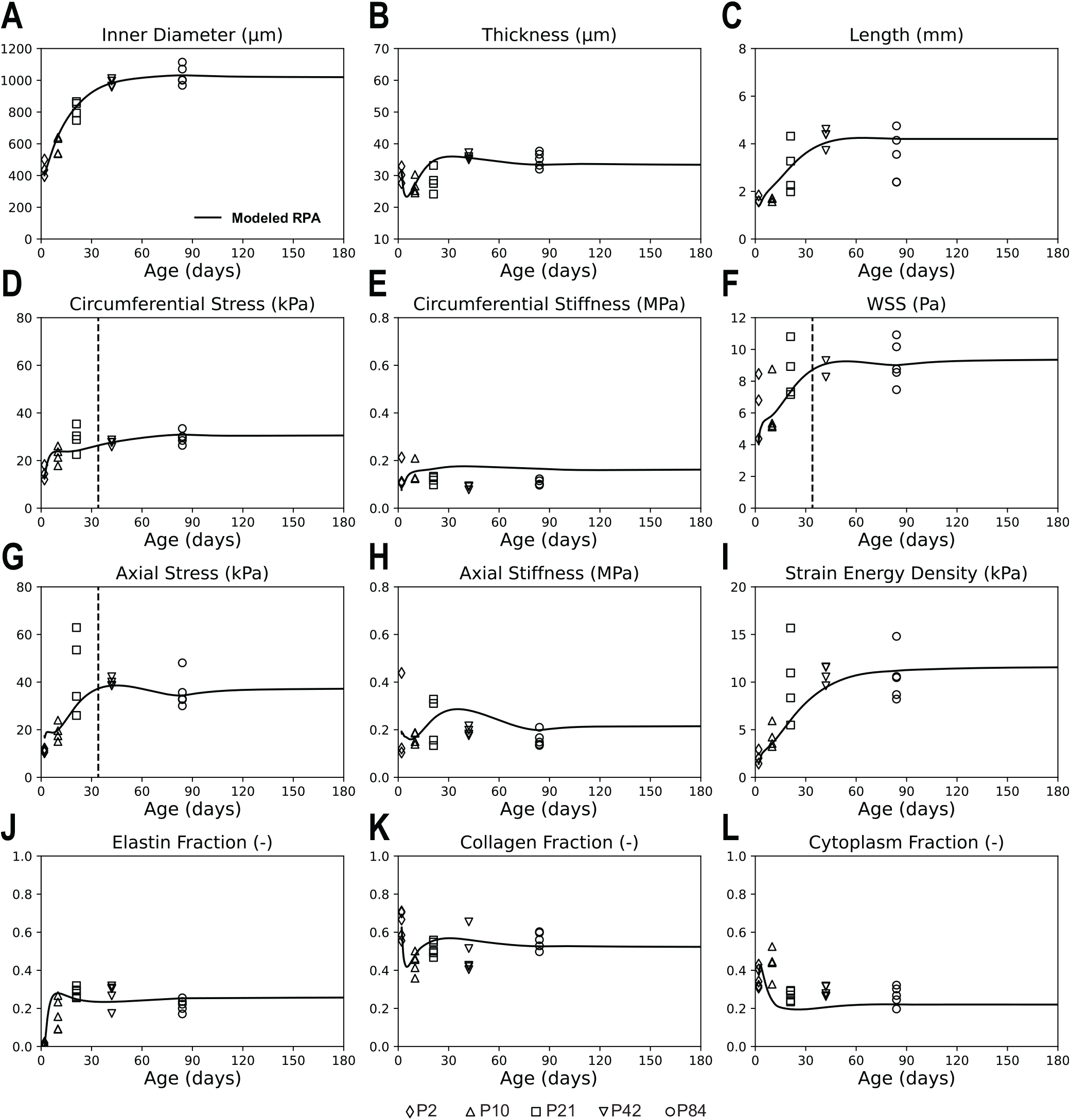
Modeled evolution of the geometry, composition, and material properties of the RPA during postnatal development (solid line) compared to observed data (symbols) at respective ages. Loaded dimensions (A, B, C), stresses (D, F, G), stiffnesses (E, H), strain energy density (I), and constituent volume fractions (J, K, L) of the modeled RPA show good agreement with observed data at 15 mmHg. Intramural stress and WSS reach 5% of their homeostatic target values at P34, demarcated by the dashed vertical line (D, F, G).

Further, the simulation framework can capture impacts of hemodynamic changes both during neonatal development and maturity. Previous studies have noted that while the LPA has similar composition to the RPA with similar stress and stiffness, it is proportionally smaller which appears to be driven by the lower flow to the left pulmonary lobe (*19*). To simulate this, we scaled the flow and axial force of our simulation proportionally to that observed between the RPA and LPA in adult mice. As noted in (*19*), the axial and circumferential stresses are similar, suggesting similar homeostatic intramural stress targets. In young rats, the RPA to LPA flow ratio is 68%:32% (*41*). If this ratio holds in early development to maturity in mice, the observed radius relationship in maturity of *a*_*LPA*_ = 0.5^1/3^*a*_*RPA*_ suggests that the RPA and LPA could experience similar WSSs in maturity. This would imply similar homeostatic targets for both intramural stress and WSS.

Using the 68%:32% flow split represents an ∼50% lower flow for the LPA. Since the axial stress in the LPA has been observed to be the same as in the RPA in maturity (*19*), if *a*_*LPA*_ = 0.5^1/3^*a*_*RPA*_ to maintain the same WSS at a 50% lower flow rate, and if ***h***_*LPA*_ = 0.5^1/3^***h***_*RPA*_ to maintain the measured circumferential stress at a lower radius (recall *a* → *ε*^1/3^*a*_***h***_ and ***h*** → *ε*^1/3^***h***_***h***_ for an *ε*-fold change in flow and *γ* = 1 change in pressure), the cross-sectional area of the LPA is *A*_*LPA*_ = 0.5^2/3^ *A*_*RPA*_ = 0.63*A*_*RPA*_. Therefore, in maturity, there is a 37% lower axial force in the LPA as compared to the RPA to enforce the same axial stress observed in the RPA in maturity. We assume that the axial force is equal to that of the RPA at birth, but is modified to reach 63% of its value at P34 such that in the LPA

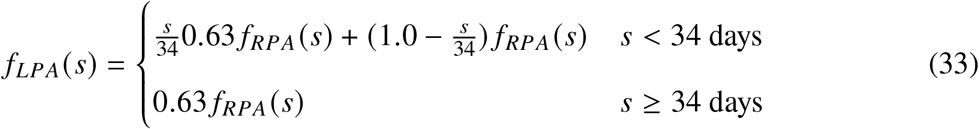

P34 is chosen as it coincides with the age at which intramural stress and WSS reach 5% of their homeostatic target values in the RPA, representing mechanical maturity (Fig. 5).

We observe that the resulting simulated LPA predicts well (Fig. 6) the measured LPA behavior in maturity (*19*). The simulated LPA has a lower inner diameter and thickness compared to the RPA, which is driven by the difference in flow. These morphologic differences arise from lower constituent family volumes, leaving the volume fraction of constituents similar to that of our modeled RPA. The stresses and stiffnesses are also similar, despite differences in morphology. This shows that our framework can capture well the normal differences resulting from hemodynamic changes superimposed on development (Fig. 6). Finally, note the long-term stability of the modeled and predicted values in maturity. This confirms both stable homeostatic responses and numerical stability over long G&R time.

**Figure 6.**
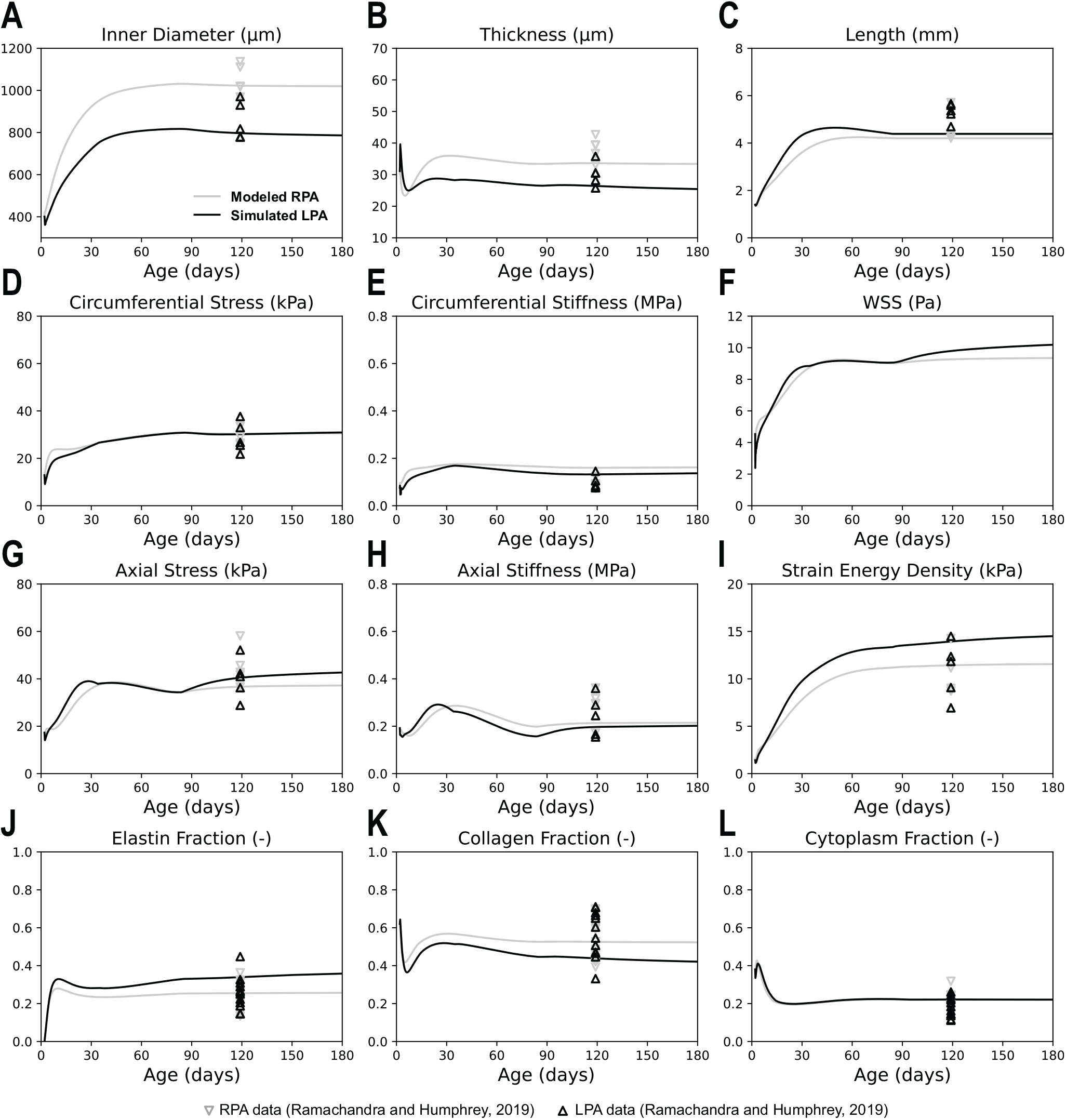
Predicted postnatal development of the LPA (dark lines) compared to the modeled development of the RPA (light lines; from Fig. 5) Symbols represent RPA and LPA data in maturity compiled from (*19*) during ages ranging from P98 to P120 but shown at P120 assuming consistent values over these ages in maturity. Inner diameter and thickness (A, B) of the LPA are proportionally smaller than that of the RPA, driven by changes in flow. Stresses (D, F, G), stiffnesses (F, H), strain energy density (I), and constituent volume fractions (J, K, L), remain similar to that of the RPA, however, which is predictive of observed LPA behavior in maturity.

This simulation framework is also capable of capturing idealized (i.e., non-immune driven, homeostatic) changes in the RPA due to hypertension in maturity (Fig. 7). Previous studies have shown varying responses of the pulmonary artery to hypertension (*46, 47*). To account for this behavior and to further parameterize the model based on experimentally measured data, we allow the mechanosensing gain-like parameters to vary in time such that

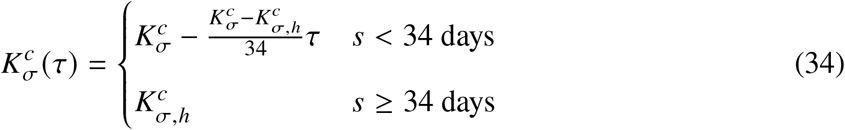

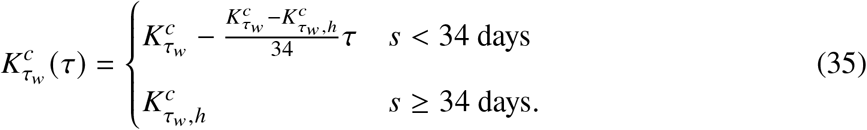

and P34 is again chosen as it coincides with the age at which intramural stress and WSS reach 5% of their homeostatic target values in the RPA, representing mechanical maturity (Fig. 5).

**Figure 7.**
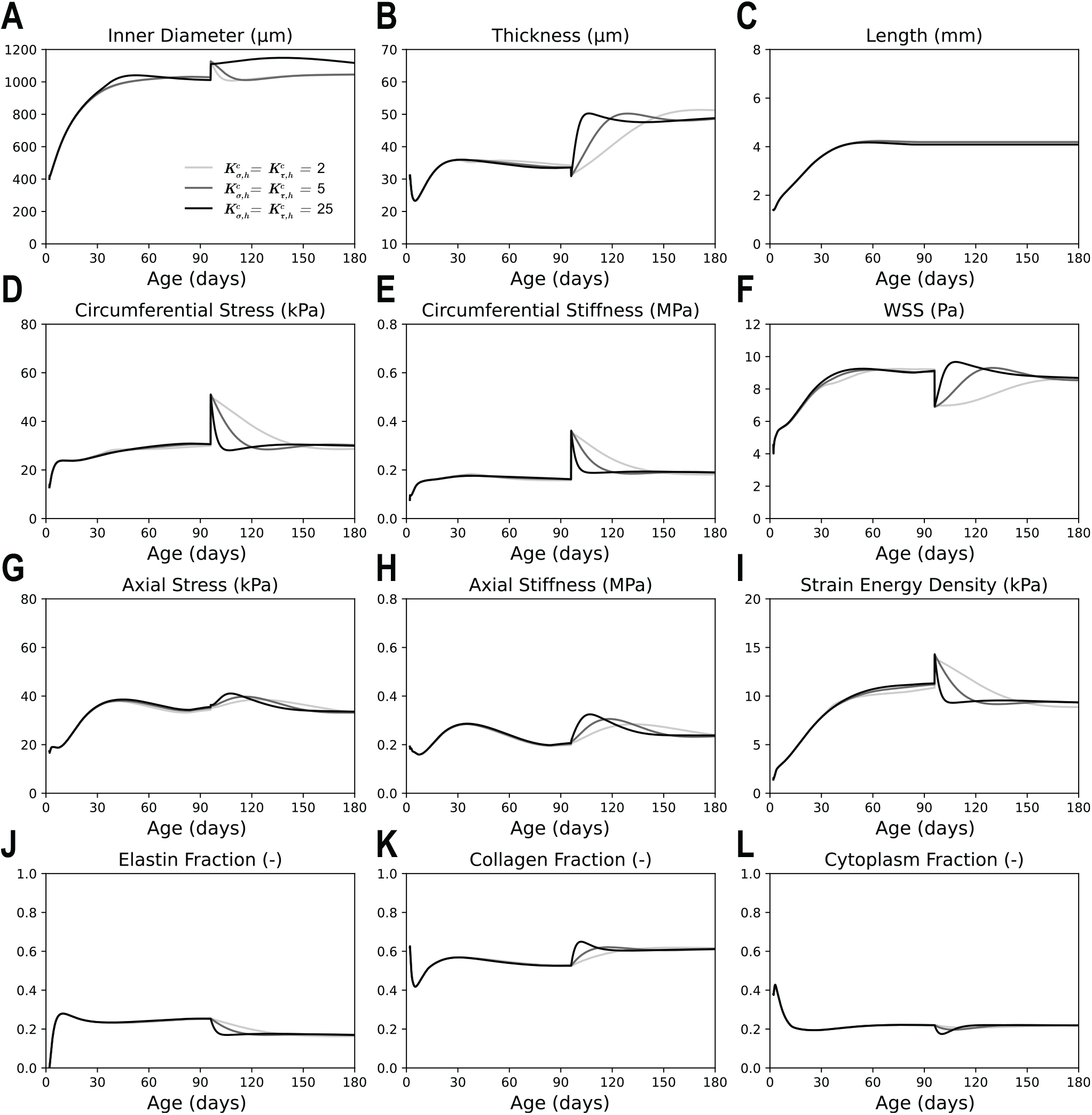
Predicted adaptation of the RPA followed by a 40% step increase in blood pressure in maturity, P96, with varying mechanosensitivity (light-to-dark lines) Diameter (A) undergoes an instantaneous increase before returning to its homeostatic value. Thickness (B) increases by approximately 40%, in line with a homeostatic response. Stresses (D, F, G) return to their homeostatic values while stiffnesses (F, H) increases slightly. Strain energy density (I) decreases slightly. Overall collagen volume fraction (L) increases during thickening, which displaces elastin volume fraction (K), while cytoplasm volume fraction (J) is mediated to return to its homeostatic density value. Rate of these changes decreases with decreasing mechanosensitivity.

Eqns. 34 and 35 hypothesize that there is high mechanosensitivity during the neonatal developmental period, which decreases linearly to the homeostatic, mature value of mechanosensitivity by P34. For all these mechanosensitivity values, pressure is increased by 40% at P96 to reach a mean arterial pressure of 21 mmHg. The vessel adapts by increasing its thickness from 34 μm to 49 μm, an approximately 40% increase in thickness consistent with the aforementioned expectation that ***h*** → *γε*^1/3^***h*** _***h***_ where *γ* = 1.4 and *ε* = 1. The rate of this increase is determined by the mechanosensitivity; higher mechanosensitivity yields a faster increase in thickness. Inner diameter, although initially perturbed by the stepwise increase in pressure, returned to its initial value to maintain WSS. The increase in thickness is primarily driven by an increase in collagen and SMCs. This results in an increase in collagen volume fraction, while the SMC volume fraction is maintained according to the density mediation outlined in Section 2.6.2. Stresses are generally maintained, but stiffnesses increase moderately. Strain energy density decreases due to the change in the elastin:collagen ratio. These simulations agree well with measured outcomes in certain cases of pulmonary hypertension, and account for the varying rates of of thickening and return to homeostatic stress targets observed in literature (*46, 47*). Importantly, these results show that the same G&R framework can capture time courses during development (Figs. 5 and 6), maturity, and disease (Fig. 7), the latter of which is capable of capturing stable adaptations to perturbation.

## 4 Discussion

Pulmonary arteries experience dramatic changes in chemo-mechanical stimuli during the perinatal period. Before birth, high values of pulmonary vascular resistance (PVR) allow the foramen ovale and ductus arteriosus to shunt blood from the right heart and proximal pulmonary artery to the left heart and systemic circulation, which is favorable because *in utero* oxygenation of blood is accomplished via the placenta. PVR decreases at birth largely due to ventilation of the lungs (*48*), with increased blood oxygenation and decreased vasoconstriction of distal pulmonary arteries also contributing (*49*). Regardless, it has long been known that natural prenatal shunting renders blood pressure nearly the same in the pulmonary artery and aorta at birth (*50, 51*), whereas pulmonary artery pressure begins to decline rapidly thereafter to near steady state values (e.g., near adult levels of ∼20/8 mmHg by 3 weeks of age in humans; (*52*)) while systemic values become progressively higher (e.g., 120/80 mmHg in adults). In mice, the foramen ovale begins to close at birth and the ductus arteriosus normally closes within hours following birth (*53, 54*), suggesting that pulmonary pressures decline more rapidly in mice and may reach near adult values by P2. Hence, in stark contrast to systemic arteries that develop postnatally under monotonically increasing blood pressure and flow (*23*), pulmonary arteries develop postnatally during a period of decreasing or stable blood pressure but increasing flow.

Based on copious studies of systemic arteries, it is generally accepted that increasing blood pressure drives wall thickness and increasing blood flow drives vascular diameter (*31, 55*). We found that wall thickness changes modestly in the RPA during postnatal development, consistent with the assumed lack of change in pressure, while diameter increases monotonically, consistent with increases in flow. Although associated developmental changes in mechanical metrics such as biaxial intramural stress, biaxial material stiffness, and stored energy appear to follow similar sigmoidal time-courses in the aorta (*23*) and pulmonary artery (Fig. 3), the magnitudes of change are very different. In the thoracic aorta, wall stress increases 4- to 9-fold, material stiffness 3- to 6-fold, and stored energy nearly 13-fold from P10 to maturity. In contrast, changes are modest in the proximal pulmonary artery where wall stress increases 1.3- to 1.9-fold, material stiffness essentially does not change, and stored energy increases 2.5-fold from P10 to maturity. We note further that blood pressure increases ∼2.5-fold in the developing aorta but not in the pulmonary artery.

Importantly, circumferential stress and material stiffness appear to be under homeostatic control in the adult aorta (*56*). Based on the prior aortic data, cell-level homeostatic values may exist early in the postnatal period whereas tissue-level values emerge only when the wall is biomechanically mature, around P56 (*23*). Findings for the RPA include potentially preferred values of cell-level circumferential stress (∼15-25 kPa) and material stiffness (∼150 kPa) from P2 to P10 but slightly higher steady state tissue-level values emerging by P34. This means, from a biomechanical perspective, that the RPA appears to mature by 4 to 5 weeks of age rather that by the 8 weeks of age needed for the aorta. If earlier maturation of the proximal pulmonary arteries in the mouse reflects a possibly analogous earlier maturation in humans, this could have consequences in early onset disease or early intervention in pulmonary care. Finally, it is interesting to compute the value of wall thickness that would be needed to increase RPA stress to aortic values (***h*** ∼4 µm, well less than a single lamellar unit) or to decrease aortic wall stress to RPA values (***h*** ∼350 µm, which would require an extensive vasa vasorum), neither of which are tenable. This comparison reveals another case of region-specific homeostasis.

Select transcriptional changes in the RPA were generally similar to those in the developing thoracic aorta (cf. (*25*)). A marker for cell proliferation (*Pcna*) decreased monotonically from P2 to maturity as did markers for the aggregating proteoglycans (*Vcan* and *Acan*) that likely facilitate the requisite cell migration while expression of genes for key extracellular matrix proteins (*Eln* as well as *Col1a1, Col3a1, Col5a1*) peaked from P10 to P21 when wall stresses were increasing most rapidly. Like values of wall stress, the associated time-course changes in histologically determined elastic fibers, fibrillar collagens, and SMCs reached steady state values by P34, seemingly consistent with the differential transcriptional profiles.

Importantly, these newly measured time-course changes in gene expression, geometry, composition, and biomechanical metrics required a new G&R framework to describe postnatal development of the RPA. In this framework, we sought to recover the principles used previously with considerable success in maturity (*30*). Key additions to the computational model included (i) presumably genetically determined developmental changes in the rates of basal mass density production and removal (initially high, but smoothly decreasing to mature values), (ii) a preferred value for SMC mass density that couples changes in cell and matrix turnover, (iii) a temporal function for collagen maturation that reflects ongoing fiber assembly and cross-linking up to mature values, (iv) a temporal function for elastic fiber maturation that reflects increasing cross-linking and compaction during early postnatal development, and (v) a possible evolution of mechano-sensitivity, as captured by temporal changes in the gain-like sensitivity parameters in the mechanical stimulus function. With these basic additions, it was possible to fit well the evolving changes in composition, geometry, and biomechanical metrics throughout postnatal development, as seen in Fig. 5. This is, to our knowledge, a novel achievement. One advantage of such a model is that data need only be collected at a small number of ages, from which the model can then interpolate for any other age within that range.

Another utility of this new G&R framework beyond describing complex evolving properties was confirmed by a simulation of LPA growth based solely on RPA growth parameters and prescription of a different evolving flowrate in the left pulmonary branch. As seen in Fig. 6, the model correctly predicted different vascular dimensions between the two branches but the same overall wall mechanics, namely wall stress, stiffness, and stored energy as reported experimentally in maturity (*19*). Moreover, the G&R framework correctly predicted ideal homeostatic adaptations in response to a sustained elevation (*γ*-fold) in blood pressure in maturity. Changes in inner radius (*a* → *ε*^1/3^*a*_***h***_) and wall thickness (***h*** → *γε*^1/3^***h***_***h***_) followed from homeostatic regulation of intramural stress and WSS as desired and demonstrated long-term stability as they should. In particular, our hypertensive model increases collagen and SMC volume (Fig. 7), resulting in arterial thickening similar to that observed in mice with induced pulmonary hypertension (*46, 47*).

Notwithstanding the success of the current approach in describing the distinctive postnatal development of the RPA, there remains a need for considerable future research. Although tissue-level homeostatic values of intramural stress emerge from the postnatal period, we were able to assume homeostatic stimulus functions throughout development because numerical differences between early and late values of stress were modest in the RPA and much of the early growth was dominated by prescribed time-courses of mass production and removal. By contrast, the early and late values of stress differ by many-fold in the aorta, and it is yet unclear if a similar approach would apply to large systemic arteries. Regardless, we do not advocate use of evolving homeostatic values during development as used before (*20, 21*) since this is inconsistent with the concept of homeostasis. Among other factors, there is a need to consider possible evolution of mechano-sensing or mechano-sensitivity, or both, particularly given the reported increase in connections between the SMCs and elastic laminae from ∼P14 to maturity in the rat aorta (*57*).

There is also a marked difference in time-course of WSS between the RPA (which increases sigmoidally and modestly) and the aorta (which experiences a transient ∼4.5-fold increase around P13 when contractile strength wanes and matrix synthesis spikes). Again, we assumed a homeostatic value for WSS throughout RPA development, which appears reasonable given the modest difference between early and late values (∼6 to ∼9 Pa) as well as prior reports that values of WSS are similar in the developing and mature pulmonary artery (*22, 41, 44*). Moreover, it is possible that our observed WSS minimum at P2 reflects a transient perturbation from its homeostatic target. Toward this end, there are several possible explanations for a transient dip in WSS during early postnatal development. Almost immediately after birth, there is a rapid increase in flow to the pulmonary arteries due to the closure of the foramen ovale and ductus arteriosus, which may cause a rapid increase in diameter that outpaces increases in cardiac output. The rapid increase in blood oxygenation at birth may also induce biochemical changes that trigger further alterations in the pulmonary arteries, including vasoactivity.

We also used the same homeostatic WSS target in the simulated LPA vessel as in the modeled RPA. Yet, (*41*) reported a higher WSS in the LPA than the RPA in rats across late postnatal ages. This may suggest different homeostatic targets for WSS in the LPA and RPA. There is evidence for this in humans as well, where WSS calculated using 4D-MRI is, by contrast, lower in the LPA than in the RPA (*58*). Given these general differences, there remains a need to identify precise WSS targets for the LPA relative to the RPA, particularly throughout development. Alternative combinations of WSS targets and flow splits could produce the observed ratio of LPA : RPA diameters in mature mice, assuming homeostatic remodeling.

Our new G&R framework nevertheless shows how changes in WSS and intramural stresses can work in tandem with mechano-mediation and homeostatic targets to yield the observed evolving pulmonary composition, geometry, and properties during development. The adversarial effects of Δ*τ*_*w*_ and Δ*σ* in Eqn. 6 ensure that both the geometry and stresses of the developing RPA remain within physiological ranges. For example, in our modeled RPA, Δ*τ*_*w*_ is slightly greater than Δ*σ* from P2 to P3.5. As deviations in WSS and intramural stress from target values modulate matrix production to similar degrees in our model (represented by equal magnitudes of the gain parameters in the stimulus function), this would cause a decrease in collagen production that associates with a reduction in thickness that aids a rapid increase in diameter during this period consistent with increased circumferential stretching. After P3.5, however, Δ*σ* becomes slightly greater than Δ*τ*_*w*_, which would cause an overall accumulation of collagen and SMCs that, in tandem with rapid collagen turnover, would restore thickness while allowing vessel diameter to increase simultaneously. By P34, both the intramural stress and WSS are within 5% of their homeostatic target, defining biomechanical maturity (Fig. 5, Fig. S3). This interaction of flow-and pressure-induced effects during postnatal development highlights the need to further understand how biomechanical perturbations drive developmental G&R. Further studies will need to examine additional biomechanical and biochemical perturbations, including complications such as inflammation in hypertension or differences due to particular pathogenetic variants.

## 5 Conclusion

It was observed nearly 50 years ago that the ascending aorta and pulmonary trunk develop very different structures due to their differing postnatal hemodynamic environments (*51*), yet relative to the extensive literature on postnatal development of the aorta (e.g., (*23, 57, 59–61*)) there has been surprisingly less attention to postnatal development of the proximal pulmonary arteries. This study successfully captured the behavior of mouse pulmonary arteries during early postnatal development and proposed a novel, data-informed G&R framework for predicting outcomes in response to hemodynamic perturbations. In this way, this study provides important insights into time-course changes in the gene expression, composition, geometry, and material properties of these arteries. While animal models will continue to provide key information, human studies will ultimately be needed to clinically translate such advances. Although compositional and structural changes in pulmonary arteries have been reported as a function of age in humans from 7 to 87 years old, the biomechanical data are often limited and do not include early postnatal periods (*62, 63*). Hence, the murine data presented here can serve as a valuable reference for future studies aimed at improving our understanding of pulmonary vascular morphogeneis, homeostasis, and pathogenesis.

## Funding

This work was supported, in part, by grants from Additional Ventures (AVCC, SVRF) and NIH (R01 HL139796, R01 HL167516). EPM is funded by the Veterans Healthcare Administration Fred Wright VISN1 CDA1.

## Author contributions

Conceptualization: ELS, ABR, DW, JDH; Methodology: ELS; Investigation: ELS, ABH, NY, EPM, DW; Visualization: ELS; Supervision: JDH; Writing—original draft: ELS, JDH; Writing—review & editing: ELS, ABR, NY, EPM, DW, JDH

## Competing interests

There are no competing interests to declare.

## Data and materials availability

The simulation framework used in this paper is available on GitHub (*64*). Data is available in the supplementary materials.

## Notes

### Competing Interest Statement

The authors have declared no competing interest.

