## Supplementary Materials for "Postnatal Pulmonary Artery Development from Transcript to Tissue"

### Supplementary Materials for
### Postnatal Pulmonary Artery Development from Transcript to Tissue

Erica L. Schwarz*, Abhay B. Ramachandra, Nicola Yeung,
Edward P. Manning, Dar Weiss, Jay D. Humphrey

**This PDF file includes:**

Figures S1 to S3

Tables S1 to S4

Reference *(64)*

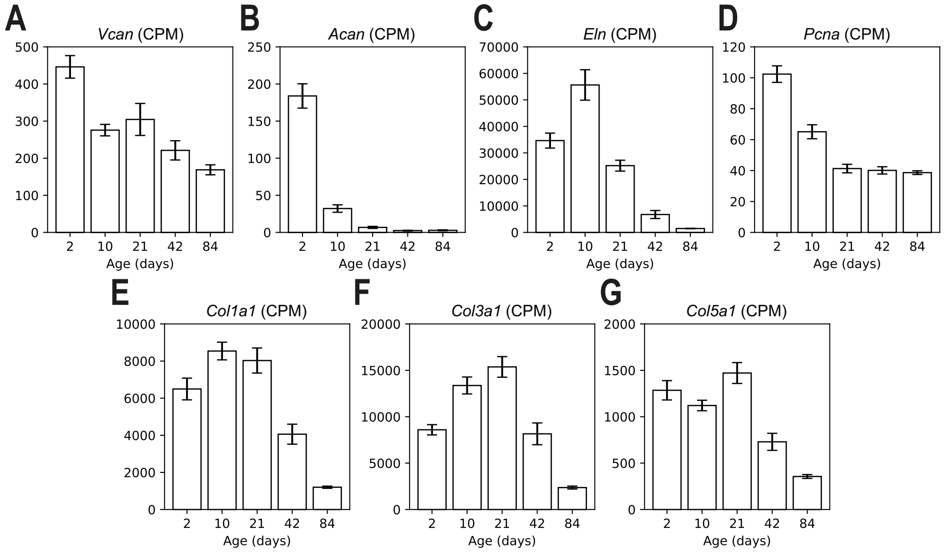


**Figure S1**: Mean ± SEM of gene expression levels in CPM (counts per million) for (A) *Vcan* (versican), (B) *Acan* (aggrecan), (C) *Eln* (elastin), (D) *Pcna* (proliferating cell nuclear antigen), (E) *Col1a1* (alpha-1 helix of collagen I), (F) *Col3a1* (alpha-1 helix of collagen III), and (G) *Col5a1* (alpha-1 helix of collagen V) during postnatal development of the RPA. Expression of *Pcna*, *Vcan*, and *Acan* decrease monotonically from P2 to lower steady state values in maturity (P84). By contrast, expression of *Eln*, *Col1a1*, *Col3a1*, and *Col5a1* peak at P10-P21, then decrease to lower steady state values in maturity.

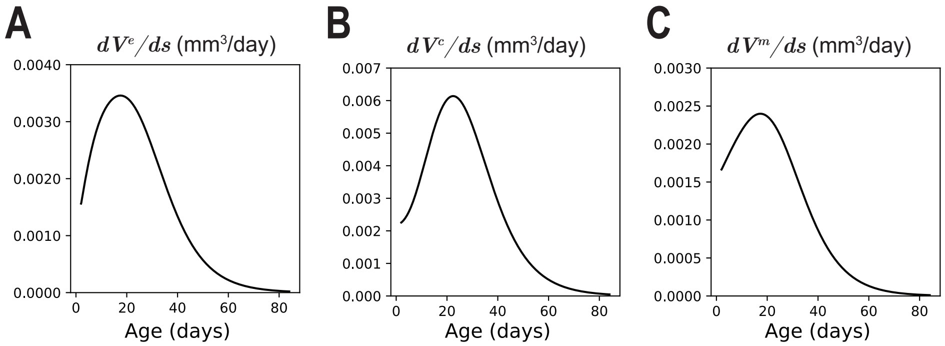


**Figure S2**: The rate of volume change of elastin (A), fibrillar collagen (B), and cytoplasm for SMCs (C).

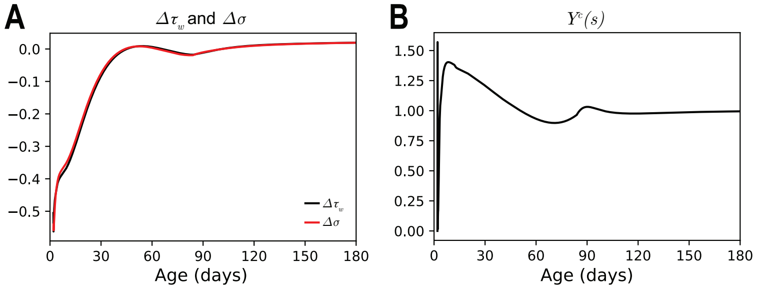


**Figure S3**: Values of $\Delta\tau_w$ and $\Delta\sigma$ (A) and stimulus function, $\Upsilon^c(\tau) = 1+K^c_\sigma\Delta\sigma(\tau)-K^c_{\tau_w}\Delta\tau_w(\tau)$, of collagen/GAGs over time (B) for the results in Fig. 5. Here, $\Upsilon^c(\tau) > 1$ represents an accumulation of collagen/GAGs, while $\Upsilon^c(\tau) < 1$ represents a loss of collagen/GAGs. Notably, $\Upsilon^c(\tau)$ is below unity from P2 to P4, likely due to reductions in the GAGs that are useful in early development when cells are proliferating and migrating. This causes a decrease in collagen production. After P4, $\Upsilon^c(\tau)$ is above unity which causes an overall increase in collagen and SMCs until reaching a stable value in maturity.

**Table S1**: Mean ± SEM of constituent area fractions calculated from Movat pentachrome (MOVAT) stained sections of the right pulmonary arteries (RPAs) at postnatal days P2, P10, P21, P42, and P84.

| | **P2**<br>**n = 5** | **P10**<br>**n = 5** | **P21**<br>**n = 5** | **P42**<br>**n = 5** | **P84**<br>**n = 5** |
|---|---|---|---|---|---|
| **Unloaded dimensions** | | | | | |
| Elastin Fraction (-) | 0.01 ± 0.00 | 0.17 ± 0.04 | 0.28 ± 0.01 | 0.27 ± 0.03 | 0.22 ± 0.01 |
| Collagen and GAG Fraction (-) | 0.64 ± 0.03 | 0.44 ± 0.02 | 0.51 ± 0.02 | 0.48 ± 0.05 | 0.56 ± 0.02 |
| Cytoplasm Fraction (-) | 0.35 ± 0.03 | 0.39 ± 0.04 | 0.20 ± 0.01 | 0.24 ± 0.02 | 0.23 ± 0.03 |

**Table S2**: Mean ± SEM morphometric and mechanical data for right pulmonary arteries (RPAs) at postnatal days P2, P10, P21, P42, and P84. The *in vivo* (loaded) values were calculated at a pressure of 15 mmHg and the individual values of axial stretch.

| | **P2**<br>**n = 3** | **P10**<br>**n = 4** | **P21**<br>**n = 4** | **P42**<br>**n = 4** | **P84**<br>**n = 5** |
|---|---|---|---|---|---|
| **Unloaded dimensions** | | | | | |
| Outer Diameter (µm) | 365 ± 11 | 405 ± 21 | 639 ± 22 | 808 ± 4 | 792 ± 18 |
| Wall Thickness (µm) | 53 ± 3.0 | 58 ± 2.0 | 61 ± 1.8 | 78 ± 1.9 | 81 ± 3.9 |
| Inner Radius (µm) | 129 ± 8 | 144 ± 11 | 258 ± 11 | 326 ± 3 | 315 ± 10 |
| Axial Length (mm) | 1.43 ± 0.04 | 1.37 ± 0.03 | 1.95 ± 0.25 | 2.74 ± 0.11 | 2.22 ± 0.27 |
| **Loaded values, Pressure = 15 mmHg** | | | | | |
| Outer Diameter (µm) | 504 ± 27 | 641 ± 27 | 872 ± 25 | 1054 ± 12 | 1101 ± 27 |
| Wall Thickness (µm) | 30 ± 1.5 | 27 ± 1.3 | 28 ± 1.9 | 36 ± 0.5 | 35 ± 1.1 |
| Inner Radius (µm) | 222 ± 15 | 294 ± 14 | 408 ± 14 | 491 ± 6 | 516 ± 13 |
| *In vivo* Circumferential Stretch ($\lambda_\theta$) | 1.52 ± 0.05 | 1.78 ± 0.06 | 1.46 ± 0.02 | 1.39 ± 0.02 | 1.50 ± 0.05 |
| *In vivo* Axial Stretch ($\lambda_z$) | 1.16 ± 0.03 | 1.22 ± 0.02 | 1.49 ± 0.08 | 1.56 ± 0.03 | 1.54 ± 0.03 |
| Circumferential Stress (kPa) | 14.86 ± 1.78 | 22.25 ± 1.78 | 29.28 ± 2.65 | 27.48 ± 0.63 | 29.60 ± 1.16 |
| Axial Stress (kPa) | 11.22 ± 0.50 | 19.01 ± 1.90 | 44.10 ± 8.53 | 40.00 ± 1.03 | 35.89 ± 3.18 |
| Circumferential Stiffness (MPa) | 0.14 ± 0.03 | 0.15 ± 0.02 | 0.12 ± 0.01 | 0.09 ± 0.01 | 0.11 ± 0.01 |
| Axial Stiffness (MPa) | 0.22 ± 0.11 | 0.17 ± 0.01 | 0.23 ± 0.05 | 0.19 ± 0.01 | 0.16 ± 0.01 |
| Strain Energy Density (kPa) | 2.13 ± 0.43 | 4.24 ± 0.60 | 10.12 ± 2.16 | 10.87 ± 0.50 | 10.56 ± 1.16 |

**Table S3**: Best-fit values of material parameters for the so-called four-fiber family model with a neo-Hookean relation capturing collective isotropic contributions to material behavior (mainly elastin) and Fung-type exponential relations capturing collective anisotropic contributions (mainly collagen fibers and passive smooth muscle).

| | Elastic fibers | Axial collagen | | Circumferential collagen + smooth muscle cell | | Diagonal collagen | | | |
|---|---|---|---|---|---|---|---|---|---|
| Age (days) | $c$ (kPa) | $c_1^1$ (kPa) | $c_2^1$ (–) | $c_1^2$ (kPa) | $c_2^2$ (–) | $c_1^{3,4}$ (kPa) | $c_2^{3,4}$ (–) | $\alpha_0$ (deg) | RMSE |
| 2 | 1.27E-12 | 1.71E-01 | 1.68E+01 | 3.91E+00 | 3.94E-14 | 1.59E-01 | 8.58E+00 | 3.24E+01 | 1.03E+00 |
| | 1.68E+00 | 6.50E+00 | 1.45E+00 | 6.98E-01 | 2.60E-10 | 1.16E-01 | 4.11E+00 | 4.45E+01 | 2.21E-01 |
| | 2.34E-14 | 1.50E+00 | 1.41E+01 | 2.69E+00 | 2.34E-14 | 8.18E+00 | 1.09E+00 | 3.22E+01 | 1.72E-01 |
| 10 | 4.10E-12 | 6.49E+00 | 3.01E+00 | 1.16E+00 | 2.36E-13 | 2.23E+00 | 8.98E-01 | 2.97E+01 | 1.41E-01 |
| | 2.37E-14 | 3.76E+00 | 3.09E+00 | 3.18E-01 | 1.86E-01 | 3.02E+00 | 8.67E-01 | 4.11E+01 | 1.27E-01 |
| | 1.31E-13 | 8.80E-02 | 4.14E+00 | 1.25E+00 | 2.83E-11 | 6.74E+00 | 3.45E-01 | 3.05E+01 | 5.80E-02 |
| | 2.34E-14 | 6.96E+00 | 6.78E-01 | 1.45E+00 | 7.38E-02 | 1.65E+00 | 1.11E+00 | 3.37E+01 | 1.35E-01 |
| 21 | 8.16E+00 | 2.64E+00 | 3.23E-13 | 4.93E+00 | 2.27E-14 | 2.79E+00 | 2.72E-01 | 3.51E+01 | 1.39E-01 |
| | 4.39E-01 | 2.01E-13 | 2.69E-14 | 4.77E+00 | 2.22E-14 | 8.28E+00 | 1.33E-01 | 3.23E+01 | 2.20E-01 |
| | 8.86E-01 | 9.05E-07 | 1.17E+01 | 2.50E+00 | 9.29E-13 | 1.26E+01 | 1.32E-01 | 3.48E+01 | 1.28E-01 |
| | 7.39E-01 | 5.65E-01 | 3.43E-13 | 2.36E+00 | 2.98E-14 | 9.87E+00 | 6.06E-02 | 3.69E+01 | 1.10E-01 |
| 42 | 9.06E+00 | 8.61E-01 | 5.82E-12 | 1.77E+00 | 2.23E-14 | 2.76E+00 | 2.64E-01 | 3.88E+01 | 1.12E-01 |
| | 8.93E+00 | 3.13E-03 | 1.61E+00 | 1.34E+00 | 8.72E-13 | 2.29E+00 | 2.60E-01 | 4.01E+01 | 8.23E-02 |
| | 1.13E+01 | 4.43E-02 | 2.20E+00 | 3.91E-01 | 7.62E-12 | 3.38E+00 | 3.28E-01 | 3.89E+01 | 8.38E-02 |
| | 8.72E+00 | 6.68E-04 | 2.85E+00 | 8.25E-01 | 2.34E-14 | 4.07E+00 | 2.58E-01 | 3.91E+01 | 1.10E-01 |
| 84 | 6.74E+00 | 2.98E+00 | 2.34E-14 | 3.67E-01 | 8.42E-13 | 1.96E+00 | 1.97E-01 | 4.03E+01 | 1.14E-01 |
| | 8.34E+00 | 1.92E-04 | 3.03E+00 | 4.72E-01 | 2.30E-10 | 2.44E+00 | 2.93E-01 | 4.20E+01 | 1.06E-01 |
| | 4.36E+00 | 1.39E-12 | 2.00E-01 | 1.57E+00 | 2.79E-14 | 5.17E+00 | 1.77E-01 | 4.06E+01 | 1.26E-01 |
| | 8.27E+00 | 1.64E-04 | 3.80E+00 | 2.83E+00 | 4.43E-14 | 3.99E+00 | 5.36E-01 | 4.09E+01 | 1.19E-01 |
| | 6.67E+00 | 1.63E-04 | 2.83E+00 | 1.05E+00 | 3.80E-14 | 3.17E+00 | 2.31E-01 | 3.76E+01 | 1.11E-01 |

**Table S4**: Simulation parameters of a developing pulmonary artery. Turnover rate parameters $a^c$, $b^c$, $k_h^c$ $a^e$, $b^e$, $a^m$, $b^m$, and $k_h^m$ are calculated from multi-modal data as specified in Section 3. The remaining parameters are fitted to capture observed evolving biomechanical and histological data of the RPA and are informed in part by parameters described previously for adult murine pulmonary arteries (*19, 45*). Note the low value of deposition stretch for elastin in the circumferential and axial directions at P2, with expected maturation of the elastic laminae between P10 and P21 (Table S3). Given that elastin is cross-linked when deposited and it does not turnover, its homeostatic pre-stretch in maturity would be expected to be between 1.1 x 1.7 (~1.9) and 1.1 x 1.25 (~1.4) based on measured developmental increases in radius (Table S2).

| Pulmonary Artery Model Parameters | | |
|---|---|---|
| Initial inner radius (mm) at Pressure = 15 mmHg | $a_0$ | 0.235 |
| Initial wall thickness (mm) at Pressure = 15 mmHg | $h_0$ | 0.027 |
| Initial length (mm) | $l_0$ | 1.4 |
| Initial volume fractions (−, −, −) | $\Phi_0^e, \Phi_0^m, \Phi_0^c$ | 0.0, 0.38, 0.62 |
| Directional collagen fractions (−, −, −) | $\beta^\theta, \beta^z, \beta^d$ | 0.37, 0.24, 0.39 |
| Initial diagonal collagen orientation (°) | $\alpha_0$ | ±30 |
| Elastin material parameters (kPa) | $c_{1_h}^e$ | 21.73 |
| Elastin deposition stretches (−, −, −) | $G_r^e$, $G_\theta^e$, $G_z^e$ | 0.1, 1.1, 1.1 |
| Elastin material maturation rates (1/days, days) | $g_1^e$, $g_2^e$ | 0.05, 21 |
| Elastin production rate constants (1/days, 1/days) | $a^e$, $b^e$ | 0.276, 0.0812 |
| Collagen material parameters (kPa, −) | $c_{1_h}^c, c_{2_h}^c$ | 155.35, 1.1 |
| Collagen deposition stretch (−) | $G^c$ | 1.21 |
| Collagen material maturation rate (1/days) | $\omega^c$ | 0.32 |
| Collagen material degradation rate (−) | $d^c$ | 8 |
| Collagen removal rate constants (1/days, 1/days, 1/days) | $a^c$, $b^c$, $k_h^c$ | 35.114, 0.0623, 0.0173 |
| Active SMC material parameters (kPa) | $T_{max_h}$ | 36.13 |
| SMC active stress remodeling time (1/days) | $k_{act}$ | 0.0169 |
| SMC baseline stretch, maximum stretch (−, −) | $\lambda_0$, $\lambda_m$ | 0.4, 1.1 |
| SMC vasoconstriction basal value, shear scaling (−, 1/Pa) | $C_b$, $C_d$ | 0.8326, 0.4163 |
| SMC production and removal rate constants (1/days, 1/days, 1/days) | $a^m$, $b^m$, $k_h^m$ | 17.533, 0.151, 0.0169 |
| Production gains (−, −, −) | $K_\sigma^c$, $K_{\tau_w}^c$, $K_\rho^m$ | 25, 25, 10 |
| Homeostatic set-points (kPa, Pa, kg/m$^3$) | $\sigma_h$, $\tau_{wh}$, $\rho_h$ | 65.47, 9.17, 231 |
